# Prediction of Hfq in actinobacteria

**DOI:** 10.1101/026195

**Authors:** Nagamani Bora, Alan C Ward, Wonyong Kim

## Abstract

Hfq is the bacterial orthologue of the eukaryotic (L)Sm family of proteins found across all domains of life and potentially an ancient protein, but it has not been found in all phyletic lines. A careful search successfully identified a distant hfq orthologue in the cyanobacteria leaving the actinobacteria as the major phylum with no known hfq orthologue. A search for hfq in actinobacteria, using domain enhanced searching (DELTA-BLAST) with cyanobacterial hfq, identified a conserved actinobacterial specific protein as remotely homologous. Structural homology modelling using profile matching to fold libraries and *ab initio* 3D structure determination supports this prediction and suggests module shuffling in the evolution of the actinobacterial hfq. Our results provide the basis to explore this prediction, and exploit it, across diverse taxa with potentially important post-transcriptional regulatory effects in virulence, antibiotic production and interactions in human microbiomes. However, the role of hfq in gram positive bacteria has remained elusive and experimental verification will be challenging.

## Introduction

Hfq is the bacterial orthologue of the eukaryotic (L)Sm family of proteins, responsible for multiple RNA processing activities^1-3^ such as RNA splicing, nuclear RNA processing and mRNA decay. The simpler hfq hexameric homopolymers provide models to understand the more complex eukaryotic pentameric heteropolymers^2^. In prokaryotes hfq^4,5^ is involved in post-transcriptional regulation^6^ acting as an RNA chaperone in facilitating interactions between small non-coding RNAs and mRNA in processes like stress responses, quorum sensing and virulence^7^.

The (L)Sm family of proteins, found across all domains of life, is potentially an ancient protein, but it has not been found in all phyletic lines^4^. A careful search successfully identified a distant hfq orthologue in the cyanobacteria^8,9^ leaving its sister clade, the actinobacteria^10,11^, with major pathogens, human commensals and industrially important species, as the major phyletic line with no hfq orthologue.

Here we show that new blast analyses of cyanobacterial hfq against actinobacterial protein sequences predict a conserved actinobacterial specific protein as distantly homologous. Secondary and tertiary structure prediction supports this with structural homology and suggests module shuffling in its evolutionary divergence. An hfq orthologue has significance for regulatory mechanisms in the corynebacteria, mycobacteria, streptomycetes and other actinobcteria.

Our results provide the basis to explore this prediction, and exploit it, across diverse taxa, in: amino acid production; nitrogen fixation; virulence; antibiotic production; and interactions in human microbiomes. Increasingly sophisticated protein structure prediction tools^12^ should uncover the roles of more small, taxon specific proteins of unknown function^13^.

## Small non-coding RNAs

Small non-coding RNAs (ncRNA) exert post-transcriptional regulation enabling rapid response to changing environmental conditions. *Cis*- antisense RNAs, transcribed from the strand opposite the target gene are complementary, and specific, to their target mRNA. *Trans*-encoded small RNAs, from intergenic regions, are weakly complementary to multiple mRNA targets, requiring hfq to facilitate interactions.

These regulatory RNAs are found across prokaryote diversity^6^. This regulatory mechanism is expected to be important in the actinobacteria. *Streptomyces* exhibit a wide range of complex regulatory processes to cope with rapidly changing environmental conditions, and diverse ncRNAs have been detected^14^. They are likely to be important tools in genetic engineering and synthetic biology^15,16^. Many *cis*- and *trans*- ncRNAs have been found in the work horse of the fine chemical industry, *Corynebacterium glutamicum*^17,18^ and their genome-wide expression demonstrated in *Mycobacterium tuberculosis*^19,20^.

## Hfq-binding small non-coding RNAs

Hfq recognises structurally diverse ncRNAs and facilitates their interaction with their partially complementary mRNA targets^1^. Hfq is the only bacterial orthologue of the (L)Sm protein superfamily. Sm proteins show a conserved Sm fold with an N-terminal α-helix followed by a twisted 5-stranded antiparallel β-pleated-sheet, with conserved motifs, Sm1 and Sm2. In hfq Sm1 is partly conserved with a different Sm2 motif, but structural studies reveal the distinct Sm fold.

A single ncRNA targeting the expression of several genes and operons can be tuned to synchronize a coordinated response to stressful environmental conditions. Such a multi-target activity requires specificity and coordinated regulation which is facilitated by hfq, which is the only factor in bacteria described as facilitating the interaction between ncRNAs and their target mRNAs^5^.

## Searching for missing Hfq orthologues

The view that blast searching might not find all hfq orthologues was supported by the failure to find hfq candidates in two major clades, the actinobacteria and cyanobacteria^4^. However, a careful search using sequence length, pattern and motifs successfully identified an ORF in the *Anabaena* PCC 7120 genome as an Hfq orthologue^8^. Blast then identified many putative homologues in unicellular and filamentous cyanobacteria and *Prochlorococcus*. The weak sequence homology was bolstered by determining the 3D structure^9^ of the *Synechocystis* sp. PCC 6803 (ssr3341 - 3HFO) and *Anabaena* PCC 7120 (asl2047 – 3HFN) hfq and their structural homology to the *Escherichia coli* hfq.

## Results

### BLAST

The actinobacteria are a sister clade to the cyanobacteria in whole organism phylogenies^10,11^. Blastp of the cyanobacterial hfq sequences against the actinobacteria found few hits with low E-values and only 2 are short (*Mycobacterium vaccae* Mvac_26751 E-value 2.1 and *Mycobacterium vanbaalenii* Mvan_1794 E-value 7.8). Blastp of the *Escherichia coli* hfq against cyanobacteria also finds few hits with E-values above the threshold, 2 are short (*Fischerella muscicola* WP_016860369.1 E-value 0.090; *Synechococcus* sp. CB0205 WP_010317051 E-value 4.1) both now annotated as hfq.

DELTA-BLAST^21^ detects remote protein homologs using domain enhanced searching and found 67 hfq matches to the *E. coli* hfq sequence in the cyanobacteria, only the last 3 are not short proteins. The top hit was the *Fischerella muscicola* hfq with an E-value of 2e-08.

DELTA-BLAST of the *Synechocystis* sp. PCC 6803 hfq against the actinobacteria matched 93 short proteins in the conserved hypothetical protein pfam family DUF3107 and aligns the DUF3107 conserved domain with the hfq and Sm-like domains in *Synechocystis* sp. PCC 6803 hfq. The top hit is to *Mycobacterium vanbaaleni* Mvan_1794 (E-value 2e-05). DELTA-BLAST of *E. coli* hfq against the actinobacteria found no hits. DELTA-BLAST of the *Synechocystis* sp. PCC 6803 hfq against the whole nr database found 2346 hits above the default threshold. The DELTA-BLAST distance tree of the results, downloaded in Newick format and displayed in Dendroscope^22^ is shown in Figure 1.

**Figure 1.**
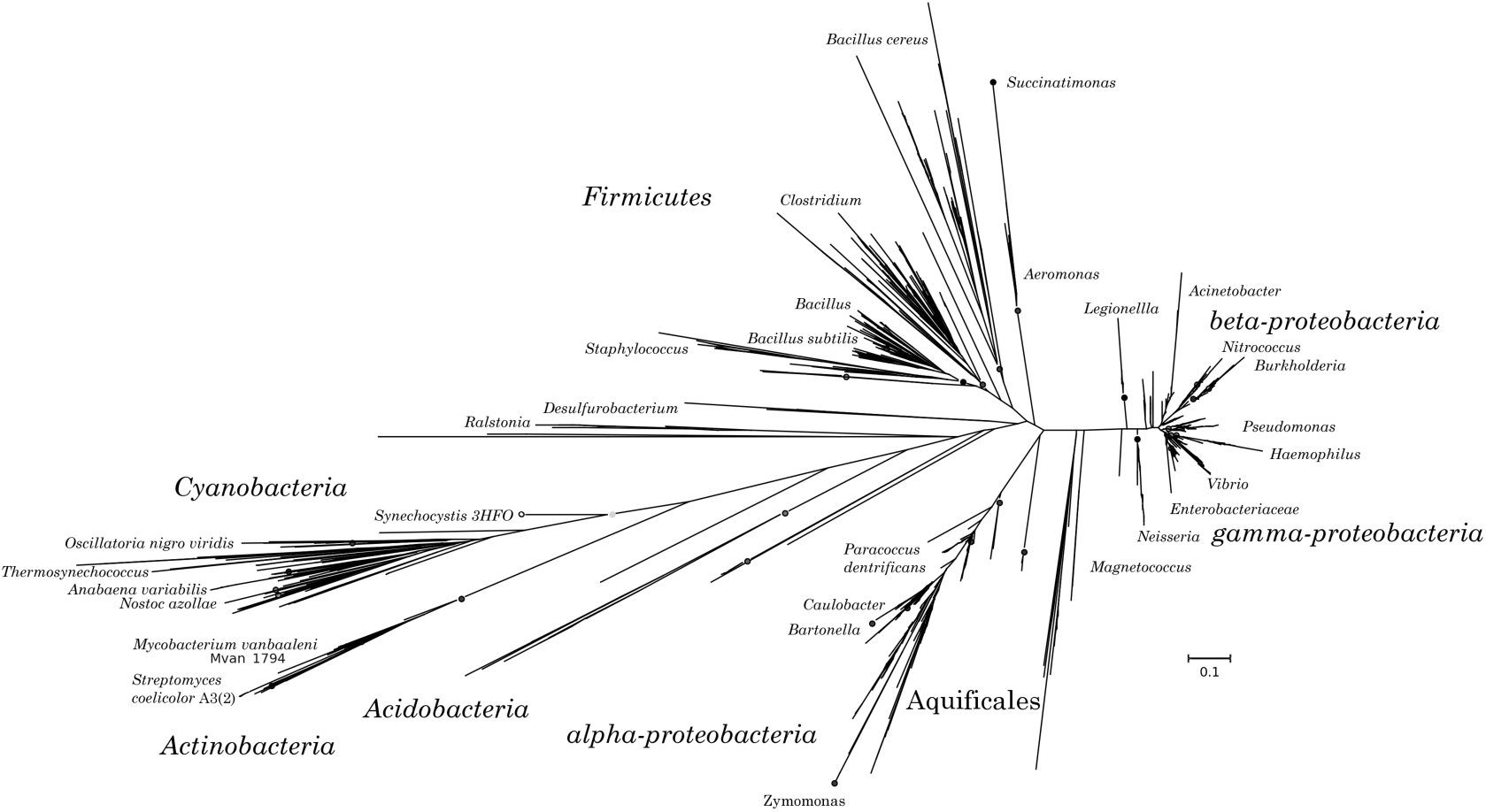
Phylogenetic tree of protein distances of significant matches in DELTA-BLAST to *Synechocystis* hfq PCC 6803 (distances for each hit are the distance from *Synechocystis* hfq)^21^.

A full, multiple sequence alignment^23^ and maximum likelihood tree^24^ displays a more distant relationship of the actinobacterial protein sequences (Supplementary data S1).

## Structural Homology

Predicting protein tertiary structure from sequence data is still challenging^12,25,26^ but *ab initio* prediction is most successful for small proteins. The Phyre2 web server^12^ generates a sequence profile of the submitted protein sequence, predicts the secondary structure (Figure 2A) and matches the profile to the profiles in a fold library.

**Figure 2.**
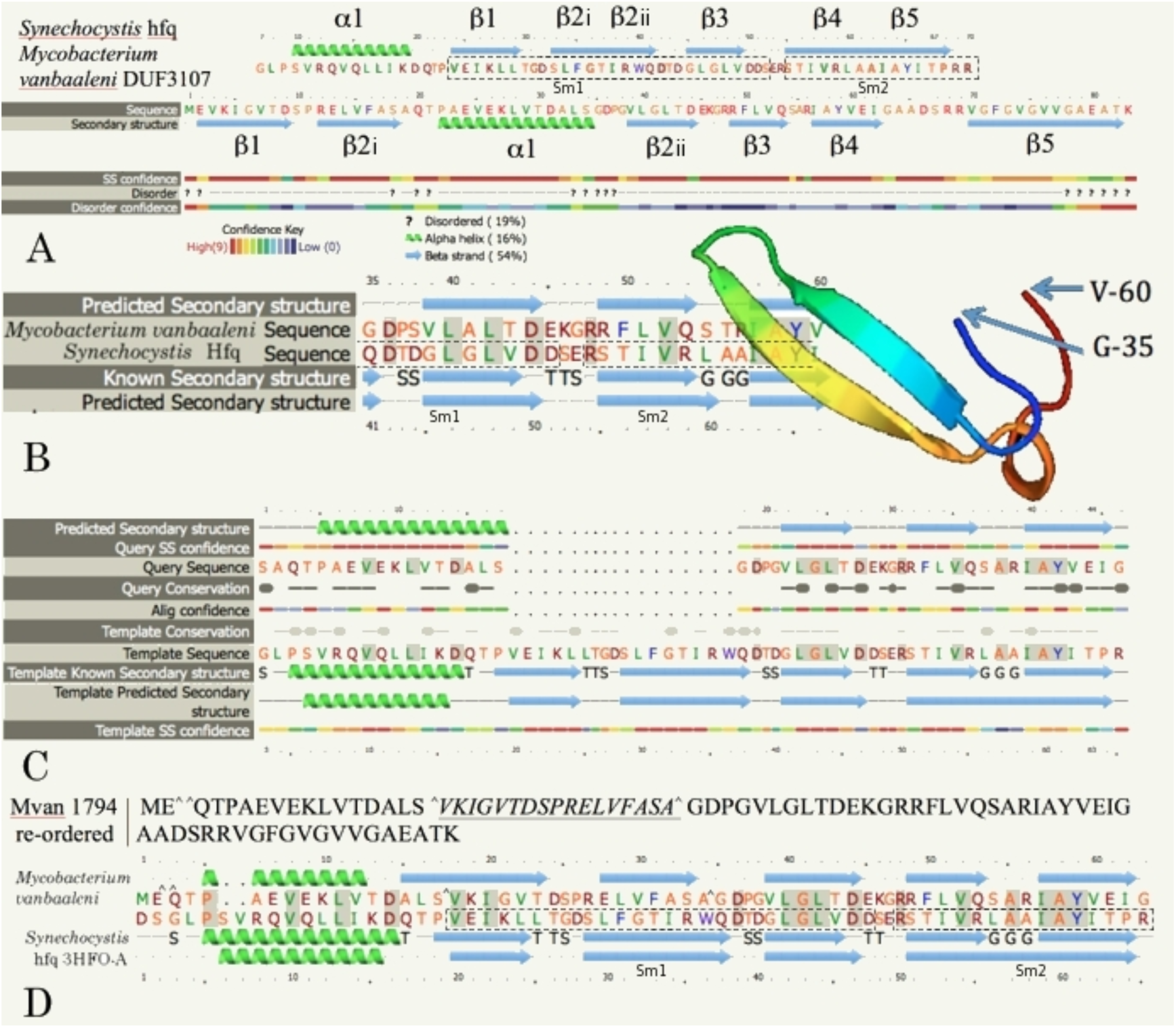
(A) Comparison of predicted secondary structure of *Synechocystis* sp. PCC 6803 Hfq. and *Mycobacterium vanbaalenii* DUF 3107 protein (Mvan_1794) from Phyre2^12^. (B) Homology modelling of the Sm-like fold in *Mycobacterium vanbaaleni* DUF3107 protein Mvan_1794 in Phyre2 to template *Synechocystis* 3HFO-C (C) One to one threading of amino acids 19-83 of *Mycobacterium vanbaaleni* Mvan_1794 against *Synechocystis* sp. PCC 6803 hfq (Phyre2 Expert mode) (D) One to one threading of the re-ordered *Mycobacterium vanbaaleni* Mvan_1794 sequence against PDB 3HFO-A, hfq from *Synechocystis* sp. PCC 6803 (secondary structure: top is actual; bottom is predicted). Mvan_1794 re-ordering: Sequence cut ^^ ^ *SEQ INSERT*^. ------ Sm1 and Sm2 in *Synechocystis* sp. PCC 6803 hfq.

The secondary structure prediction for Mvan_1794 does not show an N-terminal α-helix followed by 5-strands of β-pleated-sheet as found in hfq. However the top hit for *Mycobacterium vanbaaleni* Mvan_1794 in the Phyre2 fold library was to an Sm-like fold (Sm2) in hfq (3HFO-A) from *Synechocystis* PCC 6803 (Figure 2B). *Mycobacterium vaccae* MVAC_26751 and *Streptomyces coelicolor* A3(2) SCO5169, DUF3107 proteins with significant DELTA-BLAST matches to *Synechocystis* hfq, and *Corynebacterium glutamicum* ATCC 13032 Cgl0772 and *Bifidobacterium bifidum* ATCC 29521 BIFBIF00292, DUF3107 proteins not matched by DELTA-BLAST of the *Synechocystis* sp. PCC 6803 hfq (Supplemental data S2), returned the same hit in Phyre2.

The secondary structure of the DUF3107 sequences is 2 strands of β- pleated sheet followed by an α-helix and 4 β-pleated sheets (Figure 2A). In the 3D structure of 3HFO-A β2 is twisted, so that β2i forms an antiparallel sheet with β1, while β2ii aligns antiparallel to β3 in the β3/β4 sheet. Threading amino acids 19-83 of *Mycobacterium vanbaaleni* Mvan_1794, which includes the α-helix and succeeding β-pleated sheets, onto the template structure of *Synechocystis* 3HFO-A matches the α-helix and β-pleated sheet β3/β4, β5 of 3HFO-A (Figure 2C). The secondary structure of the DUF3107 sequences, including *Mycobacterium vanbaaleni*, and one to one threading (Figure 2) indicate that protein modules (α-helix, β1/β2i, β2ii/β3/β4, β5 and C-terminal tail) are shuffled relative to other eubacterial hfq, as β1/β2i, α1, β2ii/β3/β4, β5 and C-terminal (Figure 2D). If protein modules are identified based upon homology modelling, and the amino acids corresponding to β1/β2i in *Mycobacterium vanbaaleni* Mvan_1794 are re-ordered to follow α1, the re-ordered primary sequence threads onto the *Synechocystis* hfq sequence (Figure 2D).

The tree from the DELTA-BLAST domain-based distances (Figure 1) shows the actinobacterial sequences as a sister clade to the cyanobacterial hfq. Multiple sequence alignment and the maximum likelihood phylogenetic tree (Supplementary data S1) show the actinobacterial sequences as significantly more distant.

Linear alignment of the sequences cannot align the shuffled homologous regions, the sequence in the order β1/β2i, α1 in DUF3107 does not align with the sequence for α1, β1/β2 in 3HFO-A. The same actinobacterial primary sequences, re-ordered to match the re-arranged Mvan_1794 template (Figure 2D), align to generate a maximum likelihood tree topology (Figure 3) similar to DELTA-BLAST (Figure 1).

**Figure 3.**
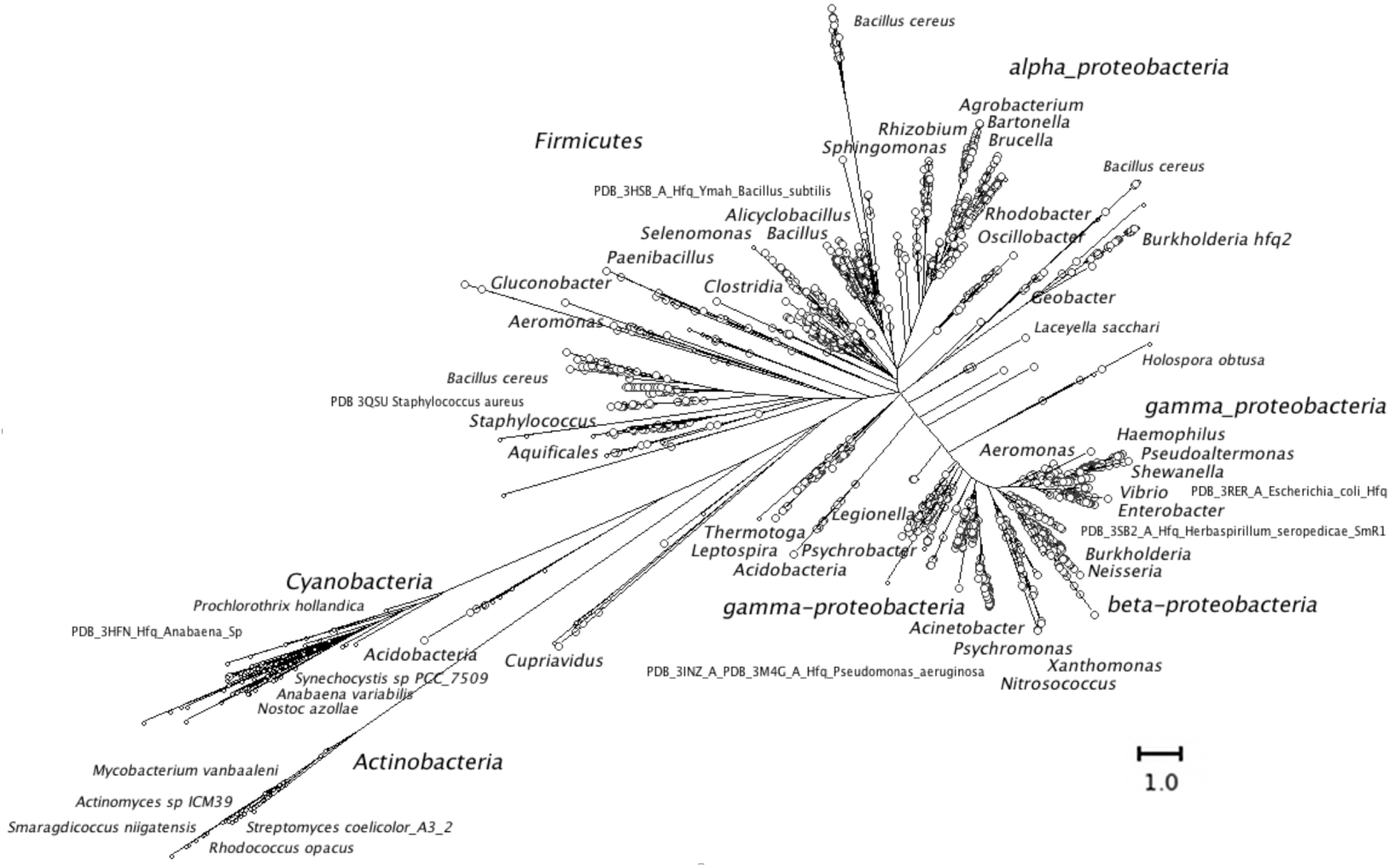
Maximum likelihood phylogenetic tree based upon multiple sequence alignment of the protein sequences retrieved using DELTA-BLAST of the *Synechocystis* sp. PCC 6803 Hfq. Aligned with the re-ordered actinobacterial sequences. ο sequences ο sequences annotated as hfq, host-factor 1 or Sm-like.

*Ab initio* modelling by Phyre2^12^ of the *Mycobacterium vanbaaleni* Mvan_1794 and *Corynebacterium glutamicum* ATCC 13032 Cgl0772 protein sequences only recovers fragments of the complete structure (Figure 2B). However, *ab initio* modelling by QUARK^26^ recovers complete 3D structure predictions, with modules that correspond to the α1, β1/β2i, β2ii/β3/β4 and β5 found in *Synechocystis* sp. PCC 6803 hfq (Figure 4 and Figure S4, supplemental data).

**Figure 4.**
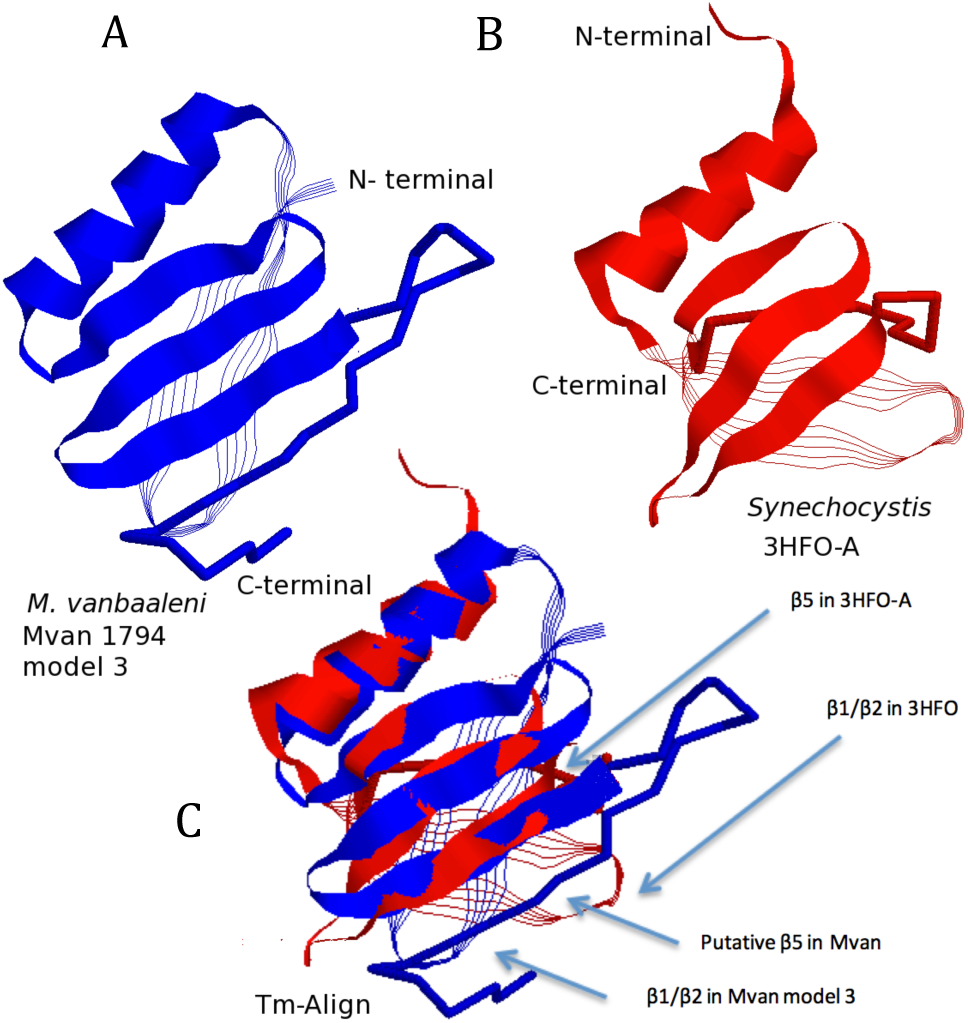
Comparison of the tertiary structures for (A) *Mycobacterium vanbaaleni* Mvan_1794 DUF3107 (Mvan_model3.pdb) (B) *Synechocystis* sp. PCC 6803 hfq (C) Aligned structures (Tm-align^27^). Aligned structures (α1 and β2ii/β3/β4) are displayed as ribbon. The β1/β2i sheet as strands and β5 in 3HFO-A, and the putative β5 and extended C-terminal in Mvan_1794, as backbone.

The best full model (Mvan_model3.pdb) aligns with the α1 and β2ii/β3/β4 modules in the *Synechocystis* sp. PCC 6803 hfq (Figure 4) with Tm-align^27^.

In Figure 4 the β1/β2i sheet of Mvan_1794 does not thread onto 3HFO-A and β5 forms antiparallel β-pleated sheet with β4 while in 3HFO-A β5 forms antiparallel β-pleated sheet with β1. The re-ordered sequence (2D) does thread the β1/β2i sheet of Mvan_1794 onto the corresponding structure in 3HFO-A (Figure S4, supplementary data).

If the predicted structure for *Mycobacterium vanbaaleni* Mvan_1794 is oriented (Figure 5A) to match the *Synechocystis* sp. PCC 6803 hfq (Figure 5D) structure, it can be assembled to match the full hexamer for 3HFO (Figure 5C and E), similar to the structure for *Salmonella enterica* hfq in^1^. In the 3HFO hexamer β5 forms antiparallel β-pleated sheet with both β1, in its own subunit, and β4, in the adjacent subunit, forming the bonds holding the hexamer together.

**Figure 5.**
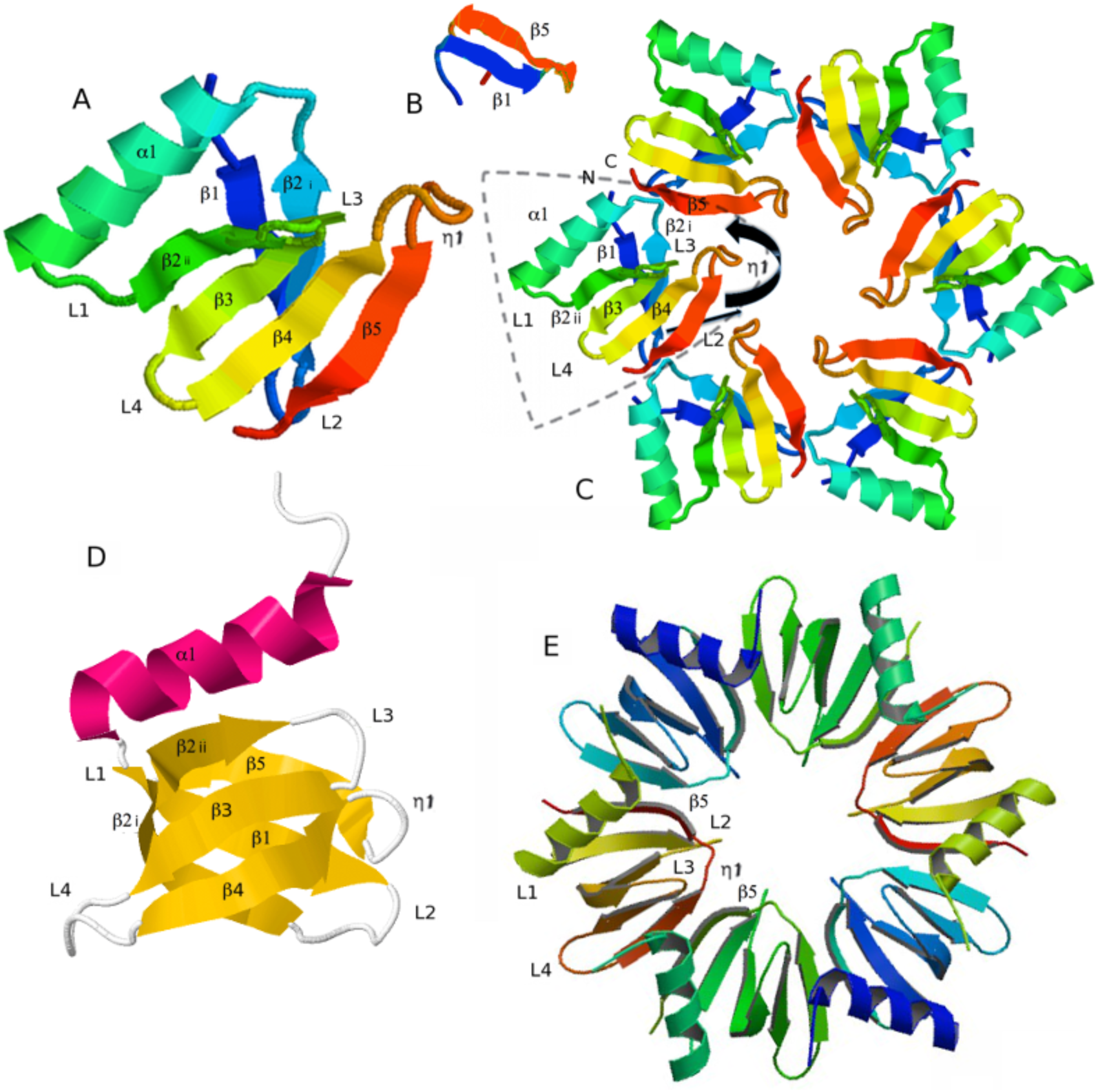
Comparison of *Mycobacterium vanbaaleni* Mvan_1794 proposed hexamer with the *Synechocystis* sp. PCC 6803 hexameric hfq. (A) predicted structure of the monomer protein of Mvan_1794 (B) - structure prediction of Mvan_1794 β1/β5 with loop η1 as linker (C) – proposed hexamer sof Mvan_1794 protein subunits. Curved arrow - proposed re-alignment of β5, straight arrow - shift of β1/β2 to interact with re-aligned β5 as modelled in (B). (D) - structure of *Synechocystis* sp. PCC 6803 hfq subunit (E) - *Synechocystis* sp. PCC 6803 hexamer.

In the model structure for *Mycobacterium vanbaaleni* Mvan_1794 the structure prediction is based upon a single polypeptide chain, and the strand proposed as the structural homologue of β5 forms anti-parallel β- pleated sheet with β4 (Figures 5A and 4). Modelling the polypeptide for *Synechocystis* sp. PCC 6803 hfq (3HFO-A) sometimes recovers the same structure in models (data not shown). In the full crystallographic structure for 3HFO β5 forms antiparallel β-pleated sheet with β1, but forms an antiparallel sheet with β4 in the adjacent subunit (Figure 4) forming the inter-subunit bonds. The β5 strand in Mvan_1794 occupies the same position as β5 strands in 3HFO, but shifted one subunit, and aligns to β4 in the same chain (Figure 5C and E). In each Mvan_1794 chain β5 could re-orientate to form β-pleated sheet with β1 (Figure 5B) and align with β4, as it currently does, but in the adjacent subunit. The sequences from β1 and β5 from the putative *Mycobacterium vanbaaleni* hfq subunit, joined with a linker polypeptide from loop η1, are modelled by Quark as β- pleated sheet (Figure 5B). This would pull β1/β2i over to move loop L2 into an even more similar position to L2 in 3HFO.

## RNA binding

The potential RNA binding residues determined from the primary sequence by BindN^29^ are shown in Figure S3 and compared with the RNA binding sites on *Salmonella enterica*^1^ and *Synechocystis* sp. PCC 6803 hfq^9^ (Supplementary data S3).

SPOT-seq^30^, template-based RNA-binding detection, was more accurate than other sequence or structure-based methods tested in^31^ and predicts hfq as an RNA-binding protein (z-scores of 18.5/18.3 for *Bacillus subtilis* and *Escherichia coli* hfq). The *Synechocystis* sp. PCC 6803 hfq and *Mycobacterium vanbaaleni* Mvan_1794, are not predicted as RNA-binding proteins but align with 3/5 of the same templates, but with lower z-scores (∼11 and 5 respectively).

## Conclusion

Bacterial whole genomes are full of small, hypothetical and often taxon specific genes, originally seen as potential artefacts, but now frequently, frustatingly, enigmatic. Of 29 signature proteins uniquely defining the actinobacteria^32^ only 2 were not annotated as hypothetical genes of unknown function. Successful searches for the function of such taxon specific hypothetical proteins e.g. Yeats *et al*. (2003)^33^, even with new tools and exponentially increasing databases, is slow.

One actinobacterial signature protein, represented by ML0814 (NP_301620) from *Mycobacterium leprae* in^32^, is a protein of unknown function containing the DUF3107 domain, present across the actinobacteria. BLAST or DELTA-BLAST of members of the DUF3107 family does not detect significant homology to non-actinobacterial proteins. A subset (93) of the DUF3107 family of proteins (>475) are identified as homologous to hfq by DELTA-BLAST of *Synechocystis* sp. PCC 6803 Hfq (Figure 1) against actinobacterial proteins.

The putative hfq orthologue in the actinobacteria has diverged in both sequence homology and module order, making identification^32,33^ as an Sm-like RNA-binding protein problematic. The determination of structural homology^9^ has been important in demonstrating the orthology and function of the cyanobacterial hfq. The phylogenetic relationship of the actinobacteria and cyanobacteria in the clade described as the “Terrabacter” by Battistuzzi *et al*.^34^ suggests the phylogeny guided search strategy used in this paper. The extent of the divergence of the putative hfq orthologue in actinobacteria, with a reorganised fold (Figure 5A and B), may indicate that more orfans may be divergent examples of functional proteins known in other taxa. But new tools and the detection of structural homology, with advances in protein structure prediction, may be needed for their identification. DELTA-BLAST looks like a useful tool for the putative identification of such partial homology. Module shuffling is common in higher eukaryotes, through mRNA isoforms with alternative exons, but is rarely documented in prokaryotes^35^.

The detection of a putative actinobacterial hfq orthologue opens new avenues for research in pathogens like mycobacteria^19^, the industrially important corynebacteria^36^ and streptomycetes^37^. However, although hfq is a key regulatory molecule in the proteobacteria^5^, its role in gram positive bacteria is less clear. An *hfq* deletion mutant of *Bacillus subtilis* showed no phenotypic changes over a wide range of growth conditions, except in survival in stationary phase in rich media^38^. It is highly conserved in all sequenced *B. subtilis* strains^38^ so survival may be a powerful selective force! Transcriptomic analysis detected changes in the levels of over 100 transcripts^38^ many linked to sporulation (although the strain used in the study was non-sporulating so these changes did not explain the stationary phase phenotype). Even a specialised role in stationary phase post-transcriptional regulation may be important in actinobacteria, for example in survival of latent pathogenic mycobacteria or secondary metabolism in actinomycetes. The *B. subtilis* hfq complemented the lack of hfq in *Salmonella* for only one hfq-dependent regulatory activity^38^. And the hfq from *Synechocystis* sp. PCC 6803 showed altered RNA binding^9^ and very limited ability to bind to *Escherichia coli* Hfq target RNAs *in vitro*. So the elusive role of DUF3107 proteins^33^ may be difficult to pin down experimentally.

## Methods

### Blast analysis

Blast and DELTA-BLAST analysis of the *Synechocystis* sp. PCC 6803 (3HFO_A) and *Anabaena* sp. PCC 7120 (3HFN_A) hfq sequences was performed at NCBI using blastp and DELTA-BLAST, with default parameters, except for increased maximum sequence hits returned. Data accessed February 2014.

### Multiple sequence alignment

Sequences were imported into SeaView^23^ and aligned using Muscle^39^. Maximum likelihood trees were generated in FastTree2^24^ with the -gamma option and visualised in Dendroscope^22^. Trees were imported into SeaView and sequences ordered to follow tree order, then manually re-aligned in SeaView following iterative tree generation and sequence ordering. All gaps inserted into variable C terminal tails were removed and matching tails manually aligned. Manual re-alignment was monitored with the cat20 gamma likelihood from FastTree2 using the -gamma option.

### Structural homology

Protein sequences were submitted to Phyre2^12^ and Quark^26^ for *ab initio* protein prediction. Models were aligned with reference structures using one to one threading in expert mode in Phyre2 and using Tm-align^27^. Structures were displayed in Rasmol 2.7.5.2 (http://rasmol.org/) and model quality assessed in PROQ2^28^.

### RNA binding

Potential RNA binding sites were identified with BindN^29^ and SPOT-seq^30^.

## Acknowledgements

The authors would like to acknowledge the powerful tools and data made available for *in silico* analysis, including DELTA-BLAST, PHYRE2^12^ and Quark^26^ and useful comments from the authors of PHYRE2^12^ and Quark^26^.

## Supplementary data

S1 Phylogenetic tree of multiple sequence alignment of sequences retrieved from DELTA-BLAST of the *Synechocystis* sp. PCC 6803 hfq NP_441518.1 sequence against the Genbank nr database.

S2 Phylogenetic tree of DUF3107 sequences Ο genera matched to *Synechocystis* sp. PCC 6803 hfq (3HFO-A) by DELTA-BLAST (Figure S2) • predicted structure in paper for *Corynebacterium glutamicum*,

S3 Two RNA binding sites on (A) Salmonella enterica hfq, (B) Synechocystis sp. 6803 hfq and (C) Mycobacterium vanbaaleni DUF3107 (amino acids 1-63) omitting putative RNA binding residues on the C-terminal tail.

S4 Alignment of the ab initio predicted structure, QUARK24, of Mycobacterium vanbaaleni protein Mvan_1794 (DUF3107) against the crystal structure of Synechocystis sp. PCC 6803 hfq (3HFO-A) by Tm- Align25 and one-to-one threading12. (A) Predicted backbone for Mvan_1794 with β1/β2i, α1, β2ii/β3/β4 segments labelled (B) corresponding structure for 3HFO-A (C) Tm-alignment of M_van_model3.pdb against 3HFO-A showing alignment of α1, β2ii, β3, β4 (compare with one-to-one threading in Figure 2D) (D) view of C along the α-helix and edge of the β2ii, β3, β4 sheet (E/F) the β1, β2i sheets, shown in grey in A and B, for Mycobacterium vanbaaleni (E) and Synechocystis (F) are in the opposite orientation, show limited sequence similarity and don’t align with Tm-Align. However, re-ordering the Mycobacterium vanbaaleni Mvan_1794 sequence segments from β1, β2i, α1, β2ii, β3, β4, β5 to α1, β1, β2, β3, β4, β5 (Figure 2D) allows one to one threading, with the β1/β2i sheet aligned, as shown in E/F

## Supplementary data S1

**Figure S1.**
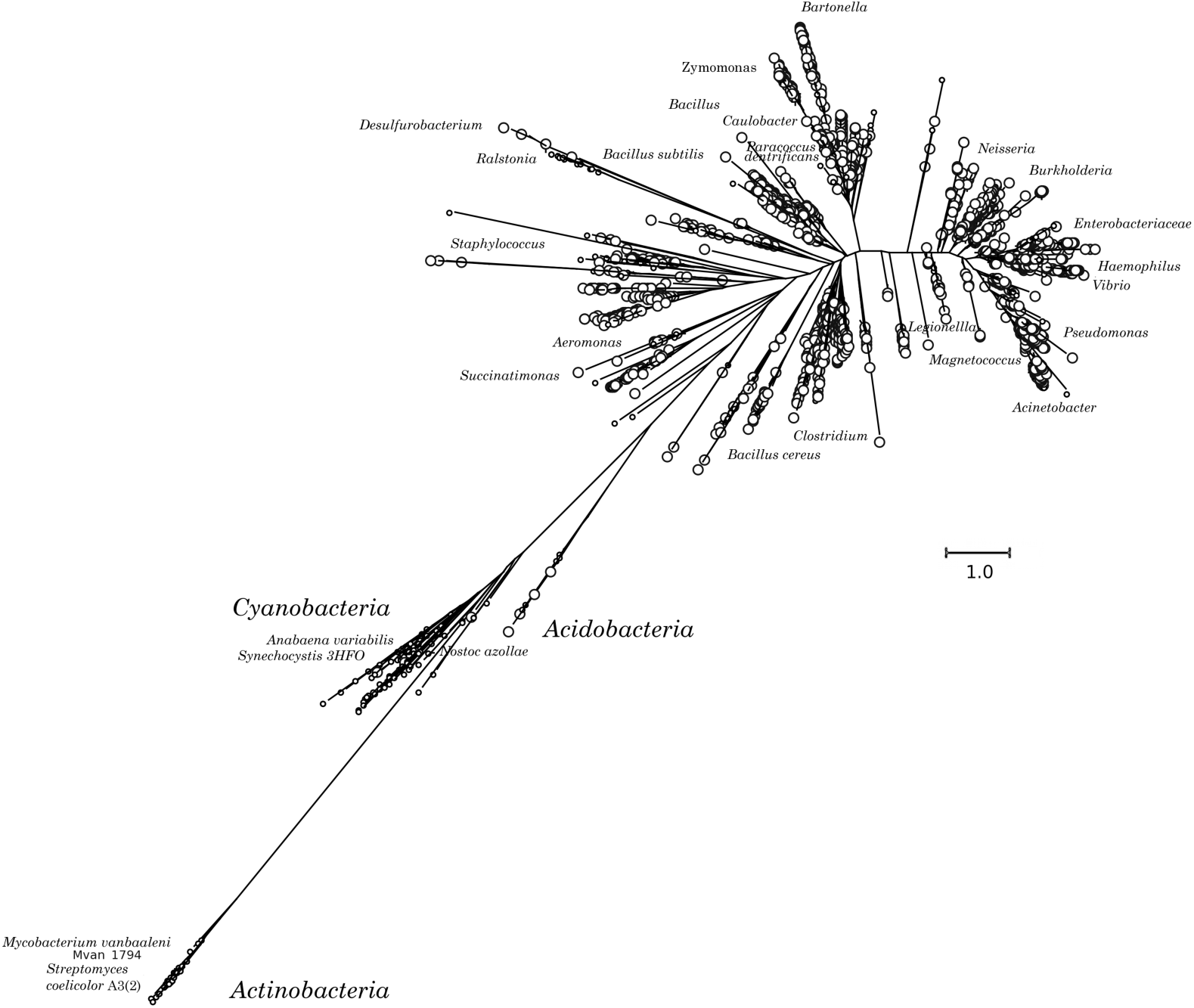
Phylogenetic tree for multiple sequence alignment of protein sequences retrieved by DELTA-BLAST of the *Synechocystis* sp. PCC 6803 Hfq. о sequences О sequences annotated as hfq, HF1 or Sm-like

## Supplementary data S2

**Figure S2.**
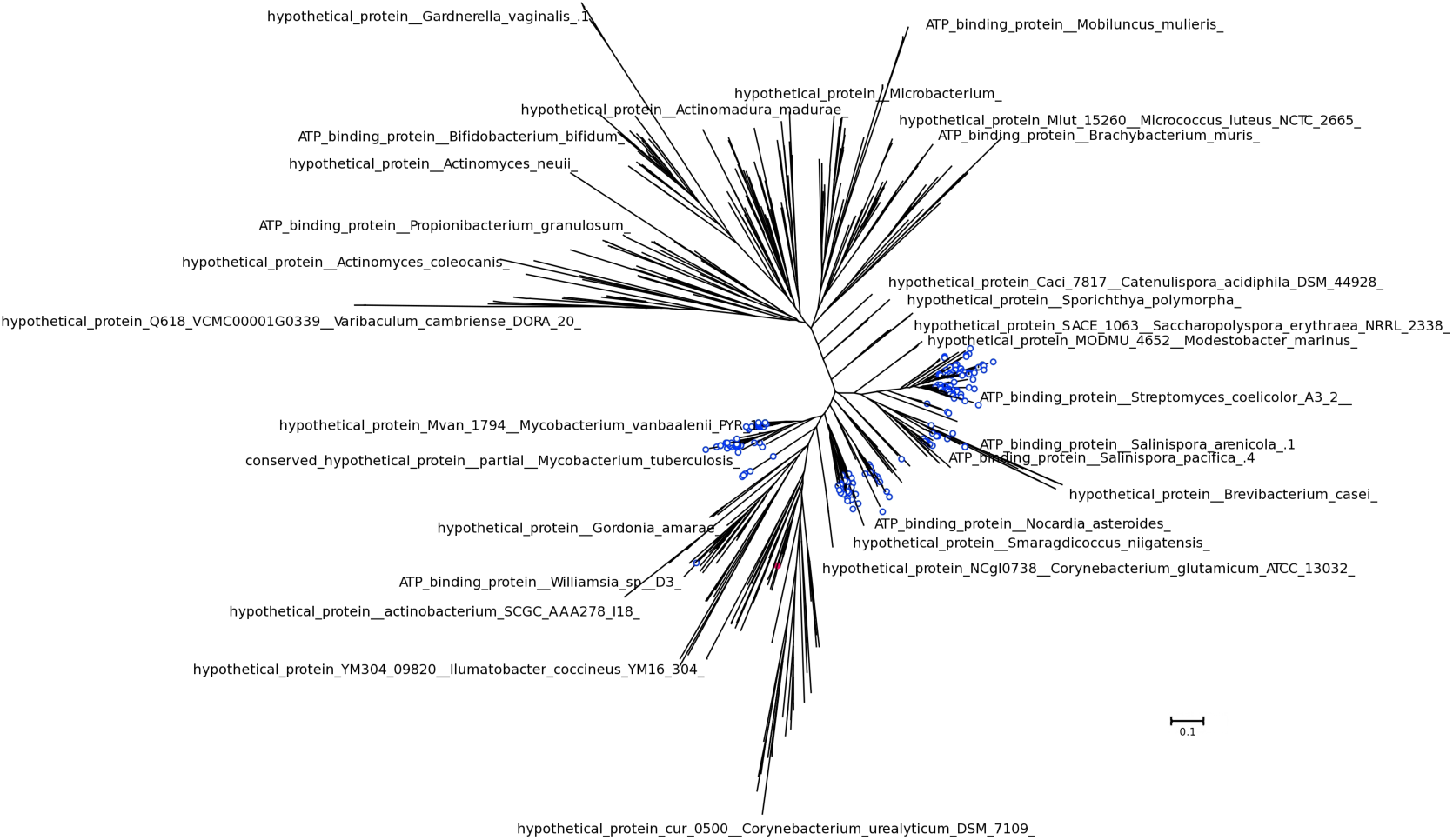
Phylogenetic tree of DUF3107 sequences • taxa matched to *Synechocystis* sp. PCC 6803 hfq (3HFO-A) by DELTA-BLAST • predicted structure for *Corynebacterium glutamicum*.

## Supplementary data S3

**Figure S3.**
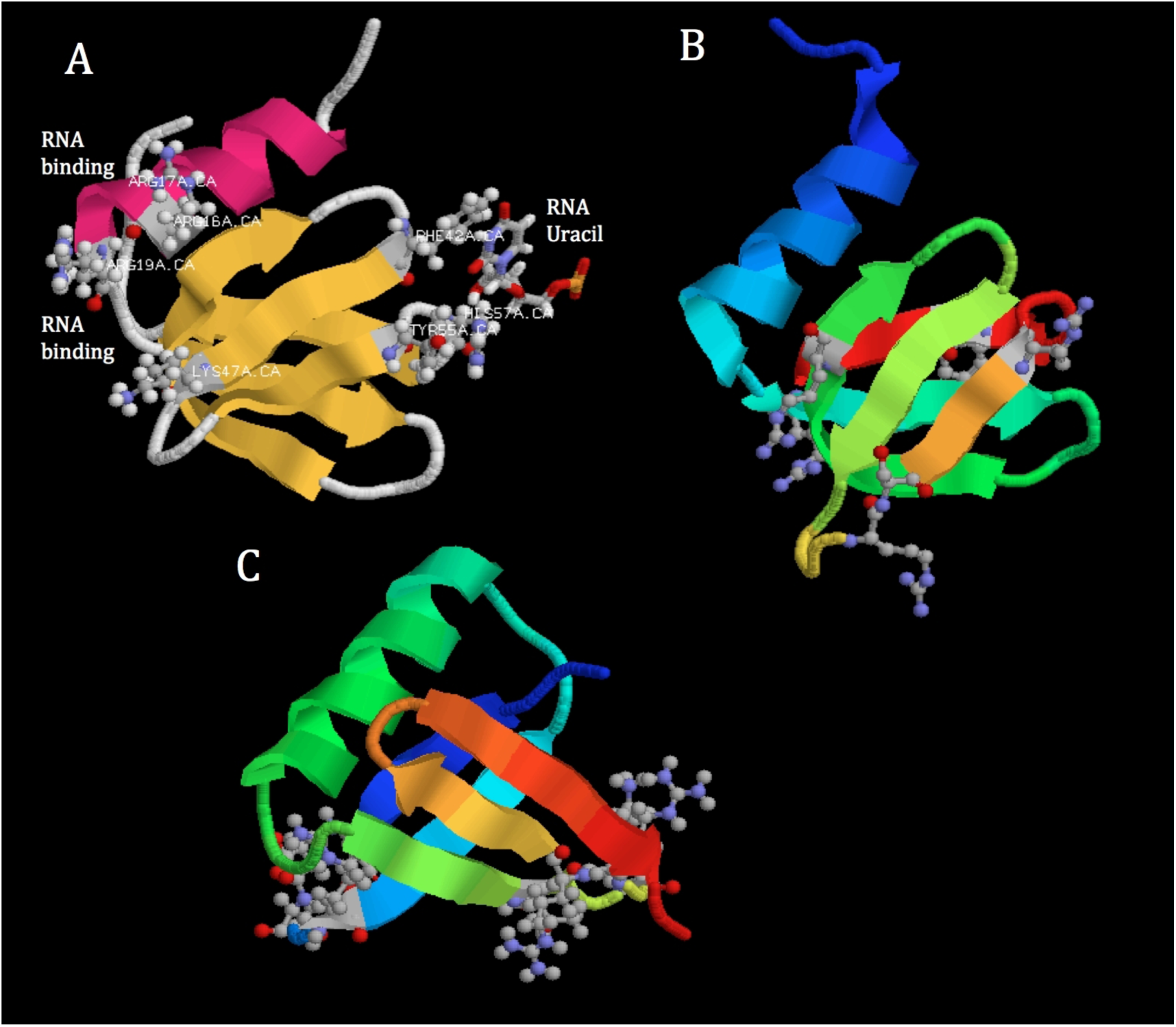
Two RNA binding sites on (A) *Salmonella enterica* hfq, (B) *Synechocystis* sp. 6803 hfq and (C) *Mycobacterium vanbaaleni* DUF3107 (amino acids 1-63) omitting putative RNA binding residues on the C-terminal tail. Supplementary data S4

## Supplementary data S4

**Figure S4.**
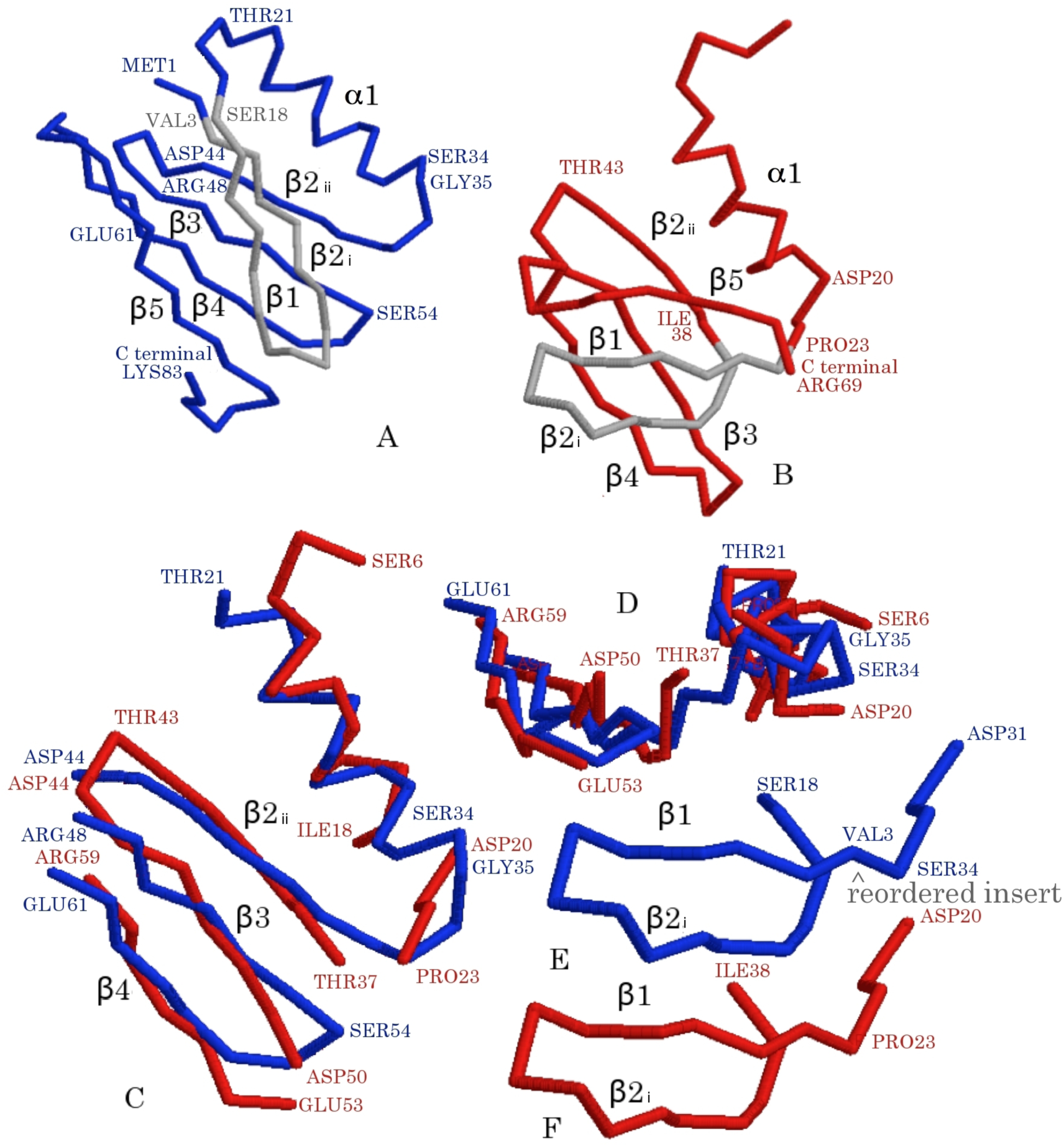
Alignment of the *ab initio* predicted structure, QUARK^26^, of *Mycobacterium vanbaaleni* protein Mvan_1794 (DUF3107 - blue) against the crystal structure of *Synechocystis* sp. PCC 6803 hfq (3HFO-A - red) by Tm-Align^27^ and one-to-one threading^12^. (A) Predicted backbone for Mvan_1794 with β1/β2i, α1, β2ii/β3/β4 segments labelled (B) corresponding structure for 3HFO-A (C) Tm-alignment of M_van_model3.pdb against 3HFO-A showing alignment of α1, β2ii, β3, β4 (compare with one-to-one threading in Figure 2D) (D) view of C along the α-helix and edge of the β2ii, β3, β4 sheet (E/F) the β1, β2i sheets, shown in grey in A and B, for *Mycobacterium vanbaaleni* (E) and *Synechocystis* (F) are in the opposite orientation, show limited sequence similarity and don’t align with Tm-Align. However, re-ordering the *Mycobacterium vanbaaleni* Mvan_1794 sequence segments from β1, β2i, α1, β2ii, β3, β4, β5 to α1, β1, β2, β3, β4, β5 (Figure 2D) allows one to one threading, with the β1/β2i sheet aligned, as shown in E/F

